# Manganese excess and deficiency affects photosynthesis and metabolism in *Marchantia polymorpha*

**DOI:** 10.1101/2022.01.24.477552

**Authors:** Marine Messant, Thaïs Hennebelle, Florence Guérard, Bertrand Gakière, Andrew Gall, Sébastien Thomine, Anja Krieger-Liszkay

## Abstract

Manganese is an essential metal for plant growth. The most important Mn-containing enzyme is the Mn_4_CaO_5_ cluster that catalyses water oxidation in Photosystem II. Mn deficiency primarily affects photosynthesis, while Mn excess is generally toxic. Mn excess and deficiency were studied in the liverwort *Marchantia polymorpha*, an emerging model ideally suited for analysis of metal stress since it accumulates rapidly toxic substances due to the absence of well-developed vascular and radicular systems and a reduced cuticle. We established growth conditions for Mn excess and deficiency, performed analysis of metal content in thalli and isolated chloroplasts and determined metabolites. Metabolome analysis revealed a strong accumulation of N-methylalanine upon exposure to Mn excess and a different response of Marchantia to heavy metal stress than that known for higher plants. We investigated photosynthetic performance by chlorophyll fluorescence at room temperature and at 77K, P700 absorption and by studying the susceptibility of thalli to photoinhibition. *In vivo* super-resolution fluorescence microscopy was used to visualize changes in the organization of the thylakoid membrane under Mn excess and deficiency. Non-optimal Mn concentrations changed the ratio of photosystem I to photosystem II and altered the organisation of thylakoid membranes. Mn deficiency seems to favour cyclic electron flow around photosystem I protecting thereby photosystem II against photoinhibition.

## Introduction

Manganese is an essential element for plant growth. Mn homeostasis is disturbed under suboptimal or excessive Mn availability (Alejandro et al., 2020). Although Mn is involved as cofactor in a range of biochemical pathways, the primary effect of Mn deficiency in photosynthetic organisms is a decrease in photosynthetic activity (Marschner, 1995). The Mn_4_CaO_5_ cluster catalyses water oxidation at Photosystem II (PSII). Furthermore, Manganese is involved in Reactive Oxygen Species (ROS) metabolism and in the antioxidant response. Mn is the cofactor of the Manganese Superoxide Dismutase (MnSOD) found in mitochondria and peroxisomes (Bowler et al., 1994; Corpas et al., 2017). Oxalate oxidase (OxOx) present in the apoplast also requires Mn. This enzyme catalyses the oxidation of oxalate to carbon dioxide coupled with the reduction of oxygen to hydrogen peroxide (Requena and Bornemann, 1999), the latter having an essential role in the defence against pathogens (Lane et al., 2002). When accumulated in excess, Mn can be toxic causing oxidative stress (Marschner, 1995; Pittman, 2005; Delhaize et al., 2007; Peiter et al., 2007; Eroglu et al., 2016).

In the presence of excess Mn in the soil, there is a competition between the uptake of Mn and other metals that are essential for the plant (Alam et al., 2005; St. Clair and Lynch, 2005; Blamey et al., 2015; Leskova et al., 2017). Indeed, at the level of the roots, plants do not have transporters that are completely selective for a single metal. Thus, high abundance of Mn can lead to a decrease in the absorption of other essential metals such as calcium, magnesium, iron or even phosphorus, negatively affecting photosynthesis (Nable et al., 1988; Amao and Ohashi, 2008) and inhibiting chlorophyll synthesis (Clairmont et al., 1986; Subrahmanyam and Rathore, 2001). Mn excess causes chlorosis followed by necrosis leading to plant death, but these symptoms are very variable and species-dependent (Millaleo et al., 2010). To overcome Mn toxicity, plants have developed different ways of Mn storage in vacuoles (Ducic and Polle, 2007), cell walls (Führs et al., 2010) or even Golgi vesicles (Marschner, 1995; Pittman, 2005). In the literature, plants have been divided into species tolerant and non-tolerant to Mn excess. Some species can hyperaccumulate Mn at levels above 10 000 mg.kg^-1^ DW (van der Ent et al., 2013).

The effect of Mn deficiency on photosynthetic electron transport and chloroplast structure has been studied for decades in a number of different organisms reaching from cyanobacteria to higher plants (e.g., Homann, 1967; Salomon and Keren, 2011). In general, a decline of photosystem II (PSII) activity is observed although symptoms of Mn deficiency are species-dependent (Homann, 1967). Depending on the severity of Mn deficiency the ultrastructure of chloroplasts may be perturbed. In slightly Mn-deficient spinach plants, the stroma lamellae are affected while the grana stacks are normal. Under more severe Mn deficiency, grana stacks are also disorganized (Mercer et al., 1962). Similar observations were reported on plants affected in Mn transporters. In Mn-deficient chloroplasts of the Arabidopsis mutant affected in Chloroplast Mn Transporter1 (CMT1), a transporter localized in the inner envelope membrane, chloroplast development was abnormal and thylakoids appeared disorganized, with either hypo- or hyper-stacked grana lamellae (Eisenhut et al., 2018; Zhang et al., 2018). In this mutant, a large heterogeneity between the chloroplasts was observed with chloroplasts containing a well-structured, normal organization of the thylakoid membrane next to chloroplasts with completely disorganized thylakoid membranes (Eisenhut et al., 2018). The consequences of the loss of PSII activity and the disorganization of the thylakoid membrane on the ratio of linear and cyclic photosynthetic electron transport have not been investigated in the previous studies.

In the present study, we investigated for the first time the consequences of both excess and deficiency of Mn on photosynthesis in the liverwort *Marchantia polymorpha*, an emerging model system. Marchantia has been described in the 1990s as being able to hyper-accumulate certain metals such as for example lead (Samecka-Cymerman et al., 1997). Due to the morphology of the thalli, the ventral part and hyaline parenchyma of the plant is in direct contact with the substrate allowing uptake of the metals from the medium without having to pass through the root system as occurs in higher plants. Compared to higher plants, the leaf anatomy of Marchantia is simpler and the tissues are thinner. Chloroplasts are less-structured than in higher plants (Tanaka et al., 2017) with smaller grana stacks, making these chloroplasts an ideal system for super-resolution fluorescence microscopy.

We have established conditions for cultivating Marchantia on plates under Mn deficiency and excess. The metal content, metabolome and antioxidant activities as well as photosynthetic activity and chloroplast structure were determined to investigate the response of Marchantia to non-optimal Mn supply. Mn excess and deficiency affect these processes in different ways. Mn excess led to a strong response of the metabolome but subtle defects in photosynthesis, while in contrast Mn deficiency affected the activity of PSII and promoted cyclic electron transport around PSI. Under Mn deficiency, an increase in non-photochemical fluorescence quenching was observed, protecting PSII against photoinhibition.

## Results

### Part I: Mn excess induces a stress response and affects the photosynthetic apparatus

To determine the effect of Mn excess on *M. polymorpha*, plants were cultivated on media containing different Mn concentrations, ranging from 33 μM (Agar Control) to 6.5 mM (200X) manganese. Gemmae were cultured on a standard Gamborg’s B5 medium (33 μM Mn) for two weeks before being transferred for one week to the different Mn concentrations before analysis. Fig. 1A shows that thalli were able to grow normally up to 1 mM Mn. However, thalli showed reduced growth from 2 mM up to 6.5 mM Mn, with signs of stress visible as brown spots. Determination of Mn concentration in thalli (Fig. 1B) showed that there is a linear correlation between Mn absorption and Mn concentration in the medium. Neither Fe nor Mg content were affected by the high Mn concentrations.

**Figure 1:**
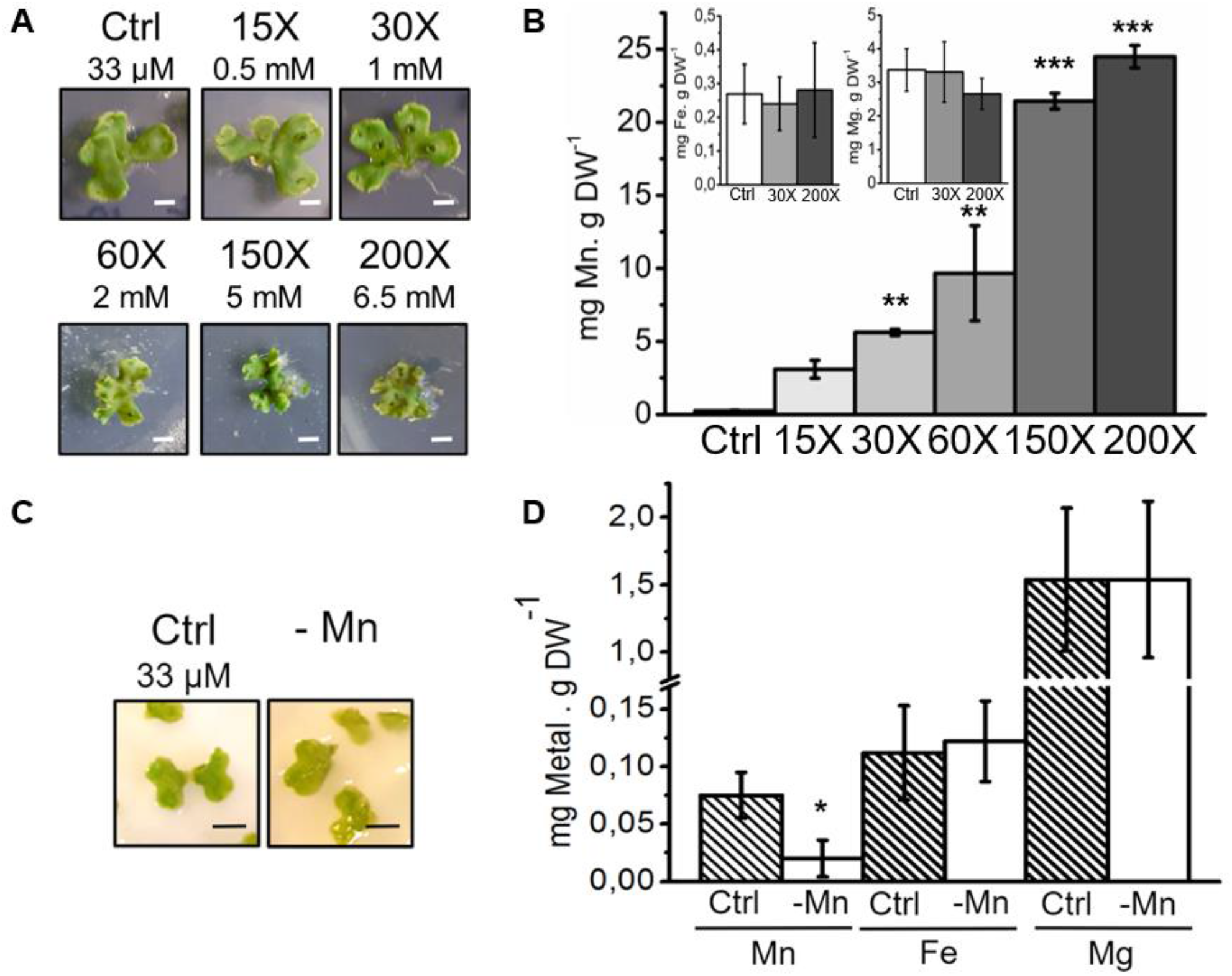
Effect of non-optimal Mn supply on growth and metal content of *Marchantia polymorpha*. **Mn excess: A:** Two-week old thalli were transferred for one week on agar plates containing 33 μM (control, Crtl), 0.5 mM (15X), 1 mM (30X), 2 mM (60X), 5 mM (150X) and 6.5 mM (200X) Mn, respectively. White scale bars represent 1 cm. **B:** Manganese, iron and magnesium concentrations were determined after one-week growth with the given Mn concentrations. **Mn deficiency: C:** Two week old thalli were transferred for one week on starch plates with or without addition of 33 μM MnCl_2_. Black scale bars represent 1 cm. **D:** Manganese, iron and magnesium concentrations in thalli were measured after one week growth on starch plates with (Ctrl, hatched bars) or without Mn (-Mn, white bars) in the medium. Error bars represent standard errors. Stars indicate significant differences, compared to the control condition, based on a Mann and Whitney test (* p<0.05, ** p<0.01, *** p<0.001; N≥3 biological replicates).

In the following, we focus on the conditions of Mn excess 30X and 200X. The first being the highest concentration at which Marchantia showed no obvious signs of stress and the second being the most stressful condition. We conducted a metabolomic analysis by GC-MS on thalli (Fig. 2). In total, 94 metabolites have been identified and quantified (SI Table S1). Under the 30X condition, there is a marked increase in threonic acid which is known to be induced in response to stress caused by heavy metals and in particular by copper in other plant species (Zhao et al., 2017). Furthermore, we observed a strong increase in fructose while a slight decrease was observed in central metabolic molecules involved in glycolysis, TCA cycle or metabolites belonging to shikimic acid derivatives compared to the control. Regarding the 200X condition, different responses occurred. Thirteen metabolites involved in the response to heavy metal or oxidative stresses were strongly increased while all other metabolites decreased compared to the control. A very high concentration of Mn is also associated with a strong induction of polyamine synthesis with an accumulation of ornithine, citrulline and putrescine. Among sugars and their derivatives, strong increases in trehalose, rhamnose and galacturonic acid are also observed. All these compounds are known to accumulate in plants and other organisms upon exposure to heavy metals (Sharma and Dietz, 2006; Hasanuzzaman et al., 2019, Liu et al., 2017; Singh et al., 2016; Alejandro et al., 2020). Most striking, N-methylalanine content was 30 and 50 times higher in 30X and 200X conditions, respectively, indicating that this molecule is the most important metabolite in the stress response induced by Mn excess in Marchantia. In parallel, the activity of a few antioxidant enzymes was measured (Table 1). A twofold increase in SOD activity was observed at the highest Mn concentration. The activity of class III peroxidases was also slightly increased at 200X, while catalase activity gradually decreased.

**Figure 2:**
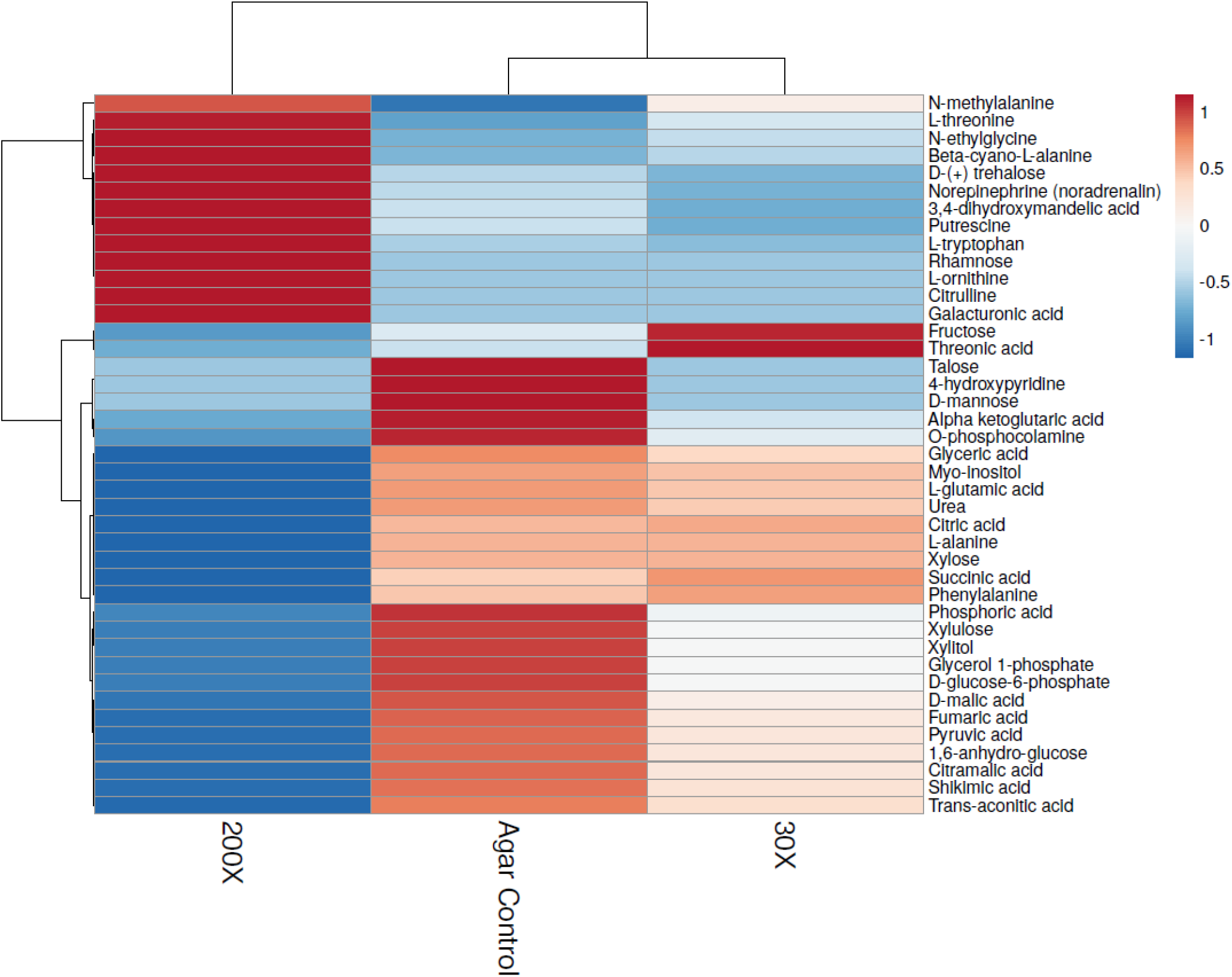
Metabolite analysis in *Marchantia polymorpha* grown in Mn excess condition. Heatmap showing significantly different metabolites in whole thalli grown under Control, 30X and 200X conditions. This figure is a result of an ANOVA 2 statistical test. The colour scale represents the variations between the three conditions taking into account the maximum and the minimum value for a given metabolite in each condition.

**Table 1.**
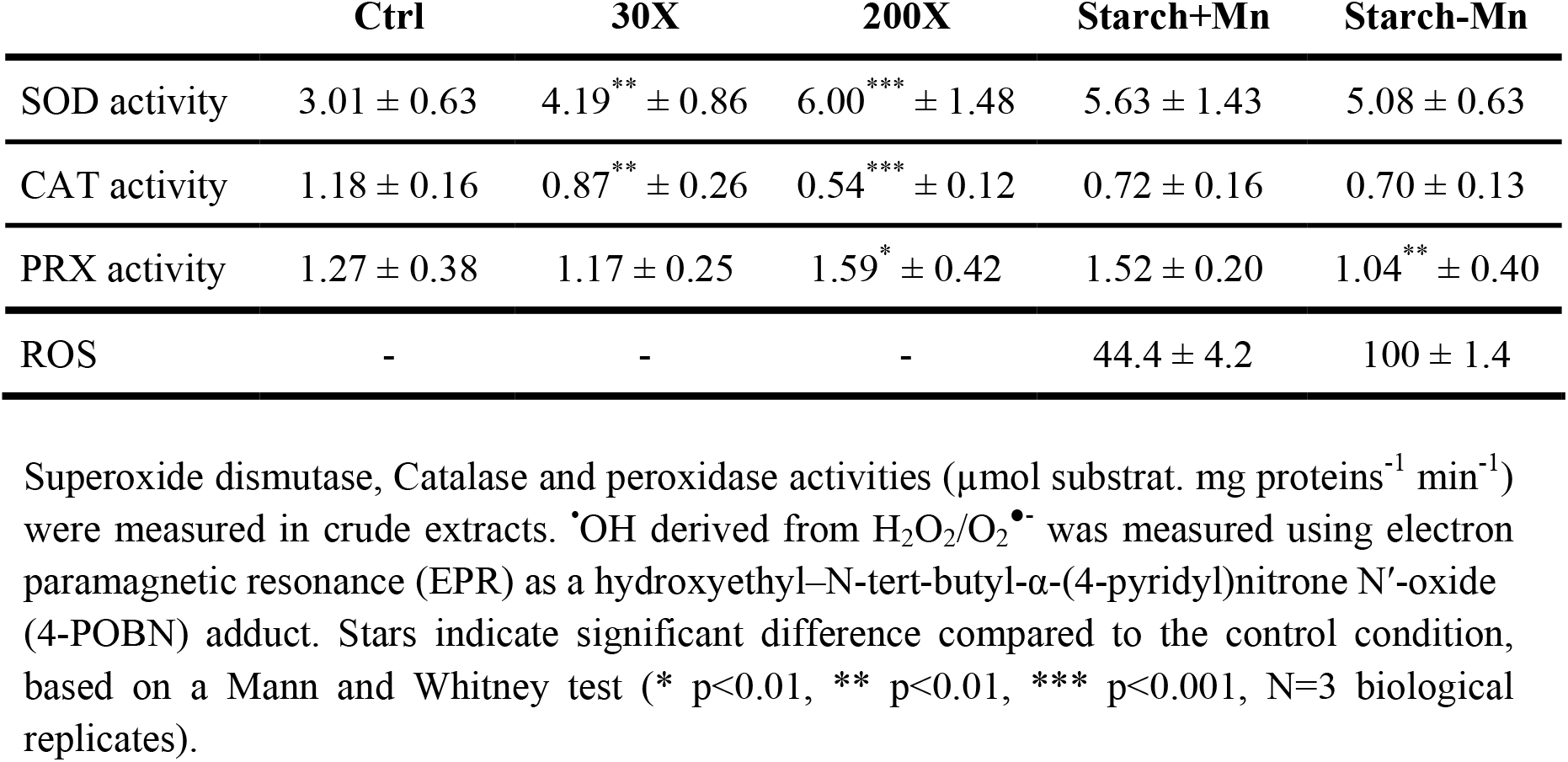
Antioxidant enzymes activities in thalli from control and manganese excess and deficiency conditions.

Next, we studied the impact of Mn excess on chloroplasts and photosynthesis. The concentration of Mn, Fe and Mg was measured in isolated intact chloroplasts (Fig. 3A). The results show a gradual and significant increase of Mn comparable with the observation made in thalli. We observed an increase from 2 to 175 μg Mn mg Chl^-1^ in 200X condition compared to the control. There were no significant changes in the Fe and Mg contents.

**Figure 3:**
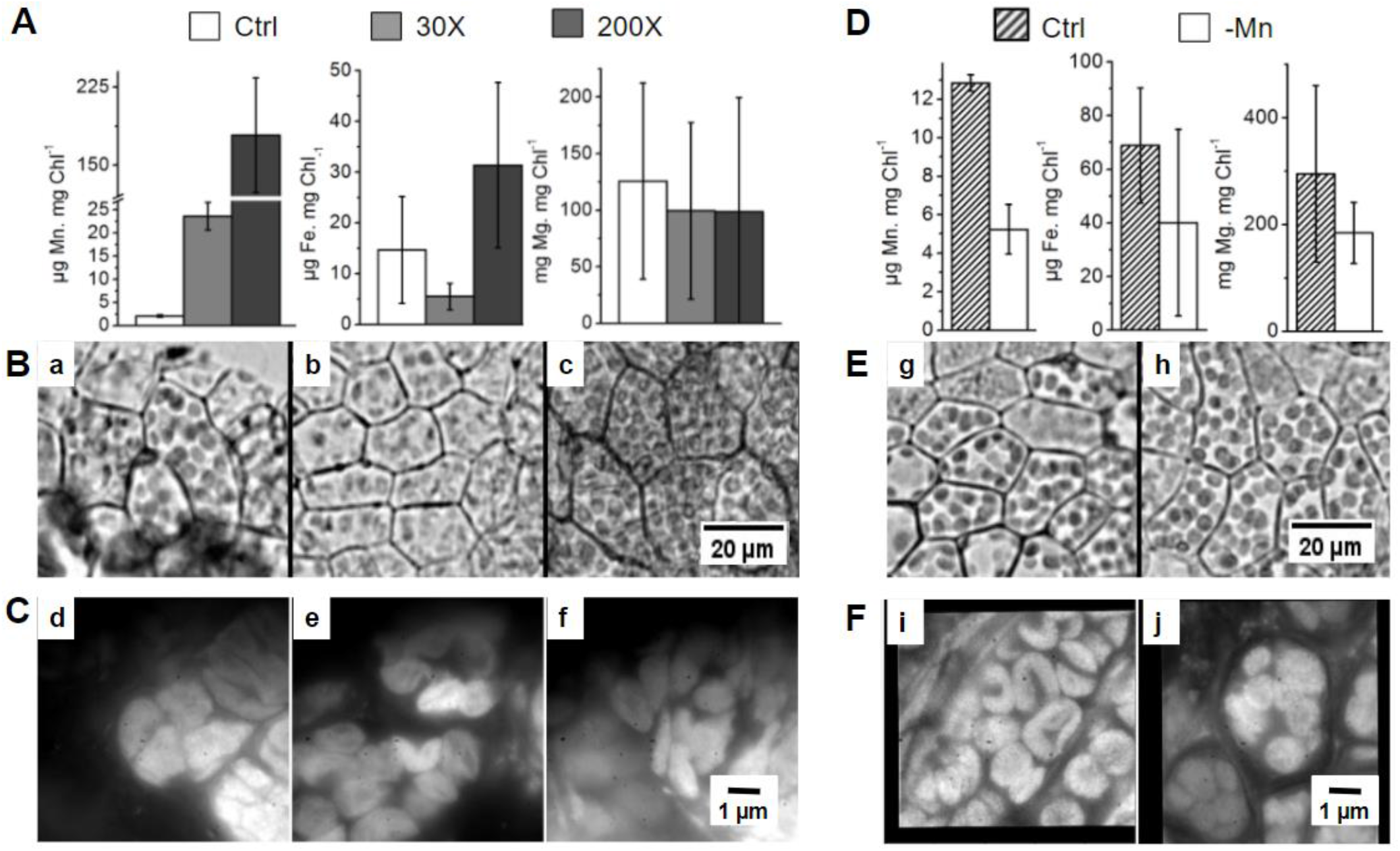
Chloroplast metal content and structure in *Marchantia polymorpha* grown on non-optimal Mn supply. **Mn excess: A:** Metal content of intact chloroplasts isolated from thalli cultured on agar (control, 33 μM; 30X, 1 mM; 200X, 6.5 mM Mn) (N=4). **B:** Microscopy images of thalli cultured on agar (a, Control; b, 30X; c, 200X). Scale bar represents 20 μm. **C:** Epifluorescence microscopy images of chloroplast surfaces of thalli (d, Control; e, 30X; f, 200X). **Mn deficiency: D:** Metal content of intact chloroplasts isolated from thalli cultured on starch (Starch+Mn, 33 μM Mn; Starch-Mn, without Mn). **E:** Microscopy images of chloroplasts in thalli (g, Starch+Mn; h, Starch-Mn). **F:** Epifluorescence microscopy images of chloroplast surfaces of thalli (i, Starch+Mn; j, Starch-Mn).

We used first bright-field and epifluorescence microscopy to see whether Mn excess changes the morphology of the cells as well as the number of chloroplasts (Fig. 3B). In Control Agar, most chloroplasts show typical plant-type spherical chloroplasts, while in 30X, there was a digression of this typical architecture. Some chloroplast had a deformed shape. In 200X, this progression of deformation of the morphology was further enhanced, and the chloroplasts appeared to be smaller on average (Fig. 3C). Next, we used super-resolution fluorescence nanoscopy to determine better the organization of the *in vivo* thylakoid membranes (Fig. 4). The organization of the chloroplast in the control (top panel) is similar to that of *Arabidopsis thaliana* (Iwai et al., 2018; Streckaite, 2021) with a round shape and the presence of numerous grana stacks seen as highly fluorescent localized fluorescence “hot-spots” (yellow/orange colour) albeit with less clear separation between stroma and grana lamellae. Under the 30X condition, the chloroplast is smaller, indicating a more compact spatial organization of the inter-connected granae. Furthermore, the presence of non-fluorescent region was observed in the centre corresponding to a void in bright field images. In 200X, the smaller chloroplasts still exhibited strong fluorescence with a well-developed distribution of spots indicative of thylakoid grana stacks.

**Figure 4:**
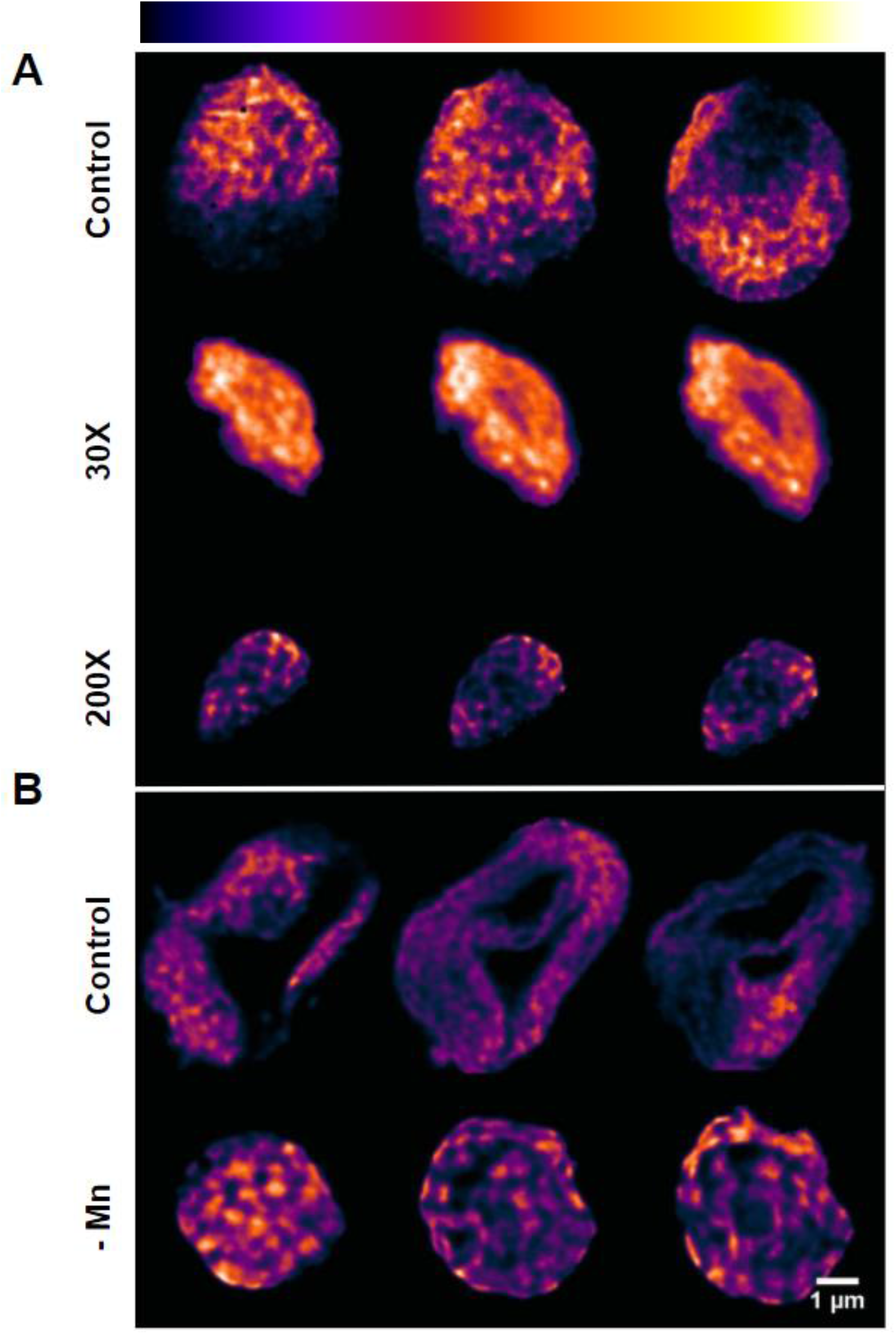
*In vivo* mesoscopic organisation of chloroplast membranes as evidenced by room-temperature super-resolution chlorophyll fluorescence emission nanoscopy. **A:** Agar Control, 30X and 200X conditions. **B:** Starch Control and Starch-Mn. For each culture condition, reading from left to right, the three reconstructed fluorescence maps are separated by 300 nm in the axial (z) direction. Light yellow represents highest fluorescence intensity and dark blue, lowest fluorescence intensity. Scale bar is 1μm. One chloroplast is represented of a total of ten chloroplasts analysed for each condition.

Since the size of the chloroplasts and the distribution of the chlorophyll fluorescence within the chloroplasts was affected by the growth conditions, photosynthesis was studied in more detail. Table 2 shows that the Chl *a*/*b* ratio decreased at 200X Mn, indicating that the antenna size relative to reaction centre increased since Chl *b* is a major constituent of the Light Harvesting Complex II (LHCII). 77K fluorescence emission spectra (Fig. 5A) showed three characteristic peaks at 685 nm, 695 nm and 735 nm, the former two peaks are attributed to LHCII and PSII, and the latter to PSI with its antenna (Satoh and Butler, 1978). For chloroplasts isolated from thalli grown at 200X, the emission attributed to PSI decreased compared to that of PSII (Fig. 5A). In line with the lower PSI fluorescence relative to PSII fluorescence in 200X, the capacity of PSI electron transport was slightly lower although the effect was not statistically significant. However, the capacity of PSII electron transfer was significantly decreased. A damaging effect of high Mn concentration on PSII activity was confirmed by fluorescence induction curves (Fig. 5B). The appearance of the K-phase in 150X and 200X conditions can be attributed to a damage of the Mn_4_CaO_5_ cluster at the PSII donor side according to Strasser (1997). In 200X, Fv/Fm and chlorophyll quenching fluorescence was altered during illumination with actinic light, but there was no increase in the susceptibility of PSII to photoinhibition (Table 2, SI Figs S1, S2).

**Table 2.**
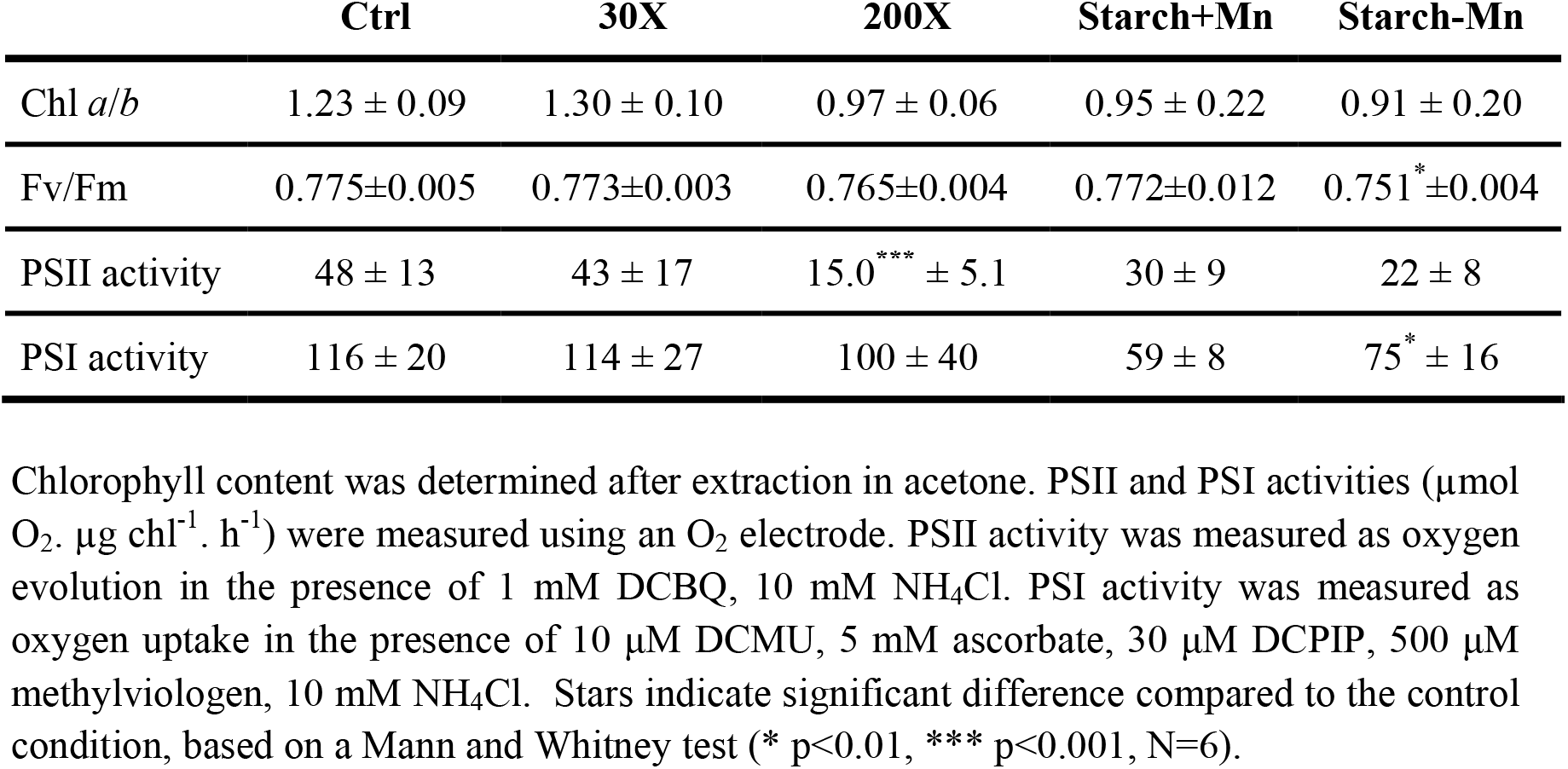
Chl *a/b* ratio and electron transport capacities of thylakoids from thalli cultured on manganese excess and deficiency.

**Figure 5:**
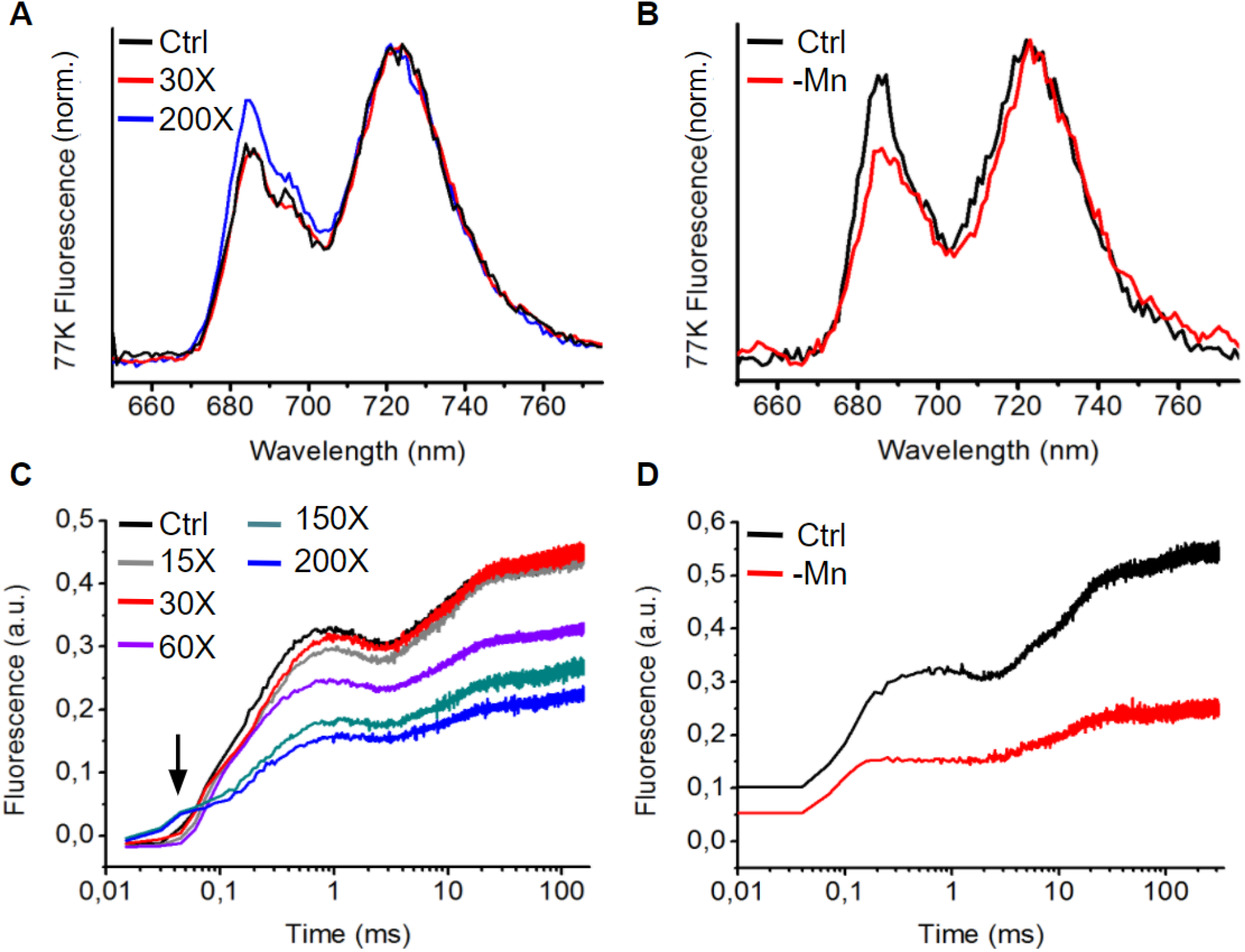
Chlorophyll fluorescence at 77K and room temperature. **A:** 77K fluorescence emission of chloroplasts extracted from thalli grown on agar: Ctrl (black), 30X (red) and 200X (blue) (N=3, biological replicates). **B:** Same as (a) carried out on starch cultivated thalli with (Ctrl, black) or without Mn (-Mn, red). **C:** Room temperature fluorescence induction curves (OJIP) were measured on 5 min dark-adapted thalli by the application of a saturating flash with 300 ms duration. Culture conditions as in **A**. **D:** Fluorescence induction curves on thalli grown as in **B**. One representative curves is shown for each condition.

### Part II: Mn deficiency affects the photosynthetic apparatus and may favour cyclic electron flow

In the following we investigated the effect of Mn starvation on *Marchantia polymorpha*. Agar plates were not suitable to induce Mn deficiency since agar contained too much Mn as impurity. Therefore, we substituted agar by starch as a gelling agent. Gemmae were cultured during 2 weeks on Gamborg’s B5 Agar medium (33 μM Mn) before young thalli were transferred on Mn-free Gamborg’s B5 Starch medium with or without addition of Mn (33 μM Mn). It was not possible to culture gemmae directly on starch because they were not able to develop. Fig. 1C shows that plants exhibited no visible growth defect after one week culturing on Starch Control or without Mn. The amount of metals in thalli was measured (Fig. 1D). The use of starch allowed a reduction of Mn content from 75 μg Mn g DW^-1^ (Starch Control) and to 20 μg Mn g DW^-1^ (Starch-Mn). Fe and Mg contents were not affected.

As for Mn excess, a metabolic analysis was carried out. No significant difference in the metabolomics profiles was observed between the two starch conditions (SI Table 1). The activity measurements of the antioxidant enzymes showed a significant decrease in the activity of type III peroxidases (Table 1). Using a spin-trapping assay, the generation of ^•^OH derived from H_2_O_2_/O_2_^•-^ was detected in starch-grown plants. ROS level were two times higher in Starch-Mn condition.

Next, the impact of the Mn deficiency on chloroplasts was investigated. The measurements (Fig. 3d) revealed a decrease in the Mn content by about 50% in the Starch-Mn condition compared with Starch Control. The Starch-Mn growth condition did not impact the iron content but slightly decreased the Mg content from 300 mg Mg mg Chl^-1^ to 200 mg Mg mg Chl^-1^. In addition, the metal content (Mn, Fe, Mg) of the chloroplasts under starch conditions was almost double compared to chloroplasts from thalli grown in the Agar Control conditions. This may be due to a higher intactness of the isolated chloroplasts. The epifluorescence microscopy images (Fig. 3f) showed more “horseshoe-shaped” chloroplasts. In Starch-Mn chloroplasts were smaller and rounder. These observations were confirmed by super-resolution nanoscopy (Fig. 4B). In Starch Control, the thylakoid membranes were organized in very distinct grana/lamellae-like structures. In Starch-Mn condition, the chloroplasts appeared to be smaller in general with fewer voids in the 3-D volume. Compared with a standard chloroplast as represented in the Agar Control (Fig. 4A, top panel), or *Arabidopsis* (Iwai et al., 2018; Streckaite, 2021), the strongly fluorescent regions were localized more distinctly, indicating a change in the mesoscopic organization of the thylakoid membrane. This may indicate smaller grana stacks with a greater segregation between grana stacks and stroma lamellae.

77K fluorescence (Fig. 5B) showed a decrease in the emission at 680 nm in chloroplasts isolated form Starch-Mn thalli, indicating a loss of the PSII content compared with PSI. PSII activity (Table 2) was lower, although the decrease is not statistically significant. Differences in the quantum yield of PSII, Fv/Fm, were significantly lower upon Mn deficiency. PSI activity increased significantly, confirming the change in the PSI/PSII ratio in Mn deficiency seen in 77K fluorescence spectra. In Starch-Mn condition, the overall fluorescence level was decreased and fluorescence induction curves showed a lower amplitude of the J phase relative to the I-P phases compared to the thalli grown on Starch Control (Fig. 5D).

Fig. 6 shows changes in room temperature chlorophyll fluorescence upon illumination with actinic light. Upon Mn deficiency, energy dissipation as heat was enhanced (qN; Fig. 6C). However, photochemical quenching (qP; Fig. 6D) was not affected. Furthermore, the slightly higher minimum fluorescence level during the recovery phase (Post-Illumination Fluorescence Transient, PIFT) upon Mn deficiency indicates an increase in chlororespiratory and/or cyclic electron flow. To show whether the stability of PSII or the repair of damaged PSII was affected under Mn deficiency, photoinhibition experiments were carried out with or without the protein synthesis inhibitor lincomycin (Fig. 7). Plants cultivated on Starch-Mn were more susceptible to high light in the absence of lincomycin. In addition, the recovery was retarded under Mn deficiency. In the presence of lincomycin, the opposite was observed with PSII in plants cultivated on Starch-Mn being more resistant to the photoinhibitory treatment. This may be explained by the higher NPQ protecting the photosynthetic apparatus.

**Figure 6:**
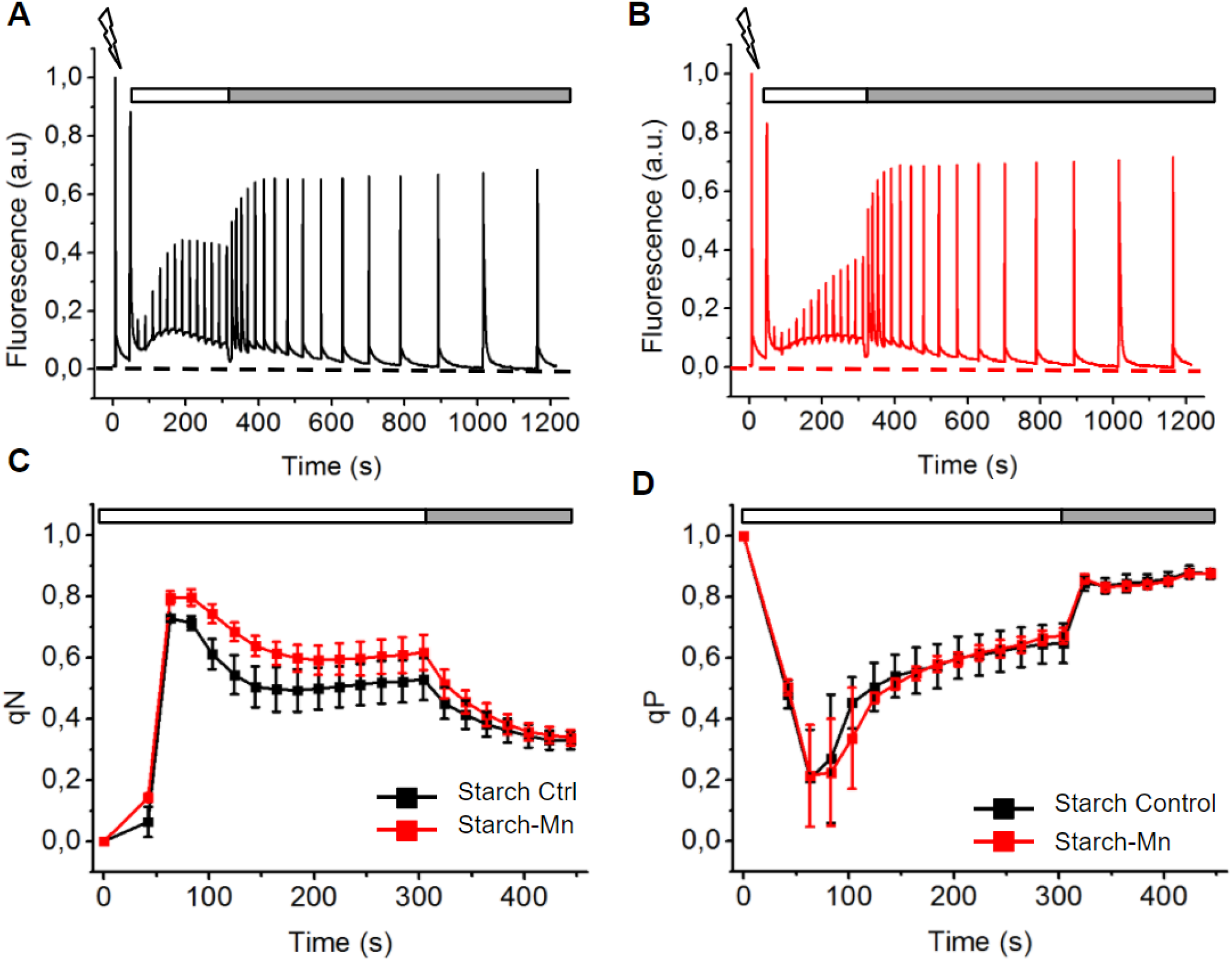
Analysis of variable chlorophyll fluorescence in Mn deficiency. **A, B:** Induction and recovery fluorescence curves were measured on thalli transferred on starch medium with (black) or without added Mn (red). After 5 min dark adaptation, thalli were exposed to a measuring light to determine the minimum fluorescence level (dashed lines) and to a saturating flash to obtain the maximum fluorescence level. Actinic light (50 μmol photons m^-2^ s^-1^) was applied for 5 min (white bar). Recovery in the dark was followed (grey bar). **C:** qN and **D:** qP parameters were calculated thanks to saturating flashes applied during the light and dark period (N=12, 3 biological replicates).

**Figure 7:**
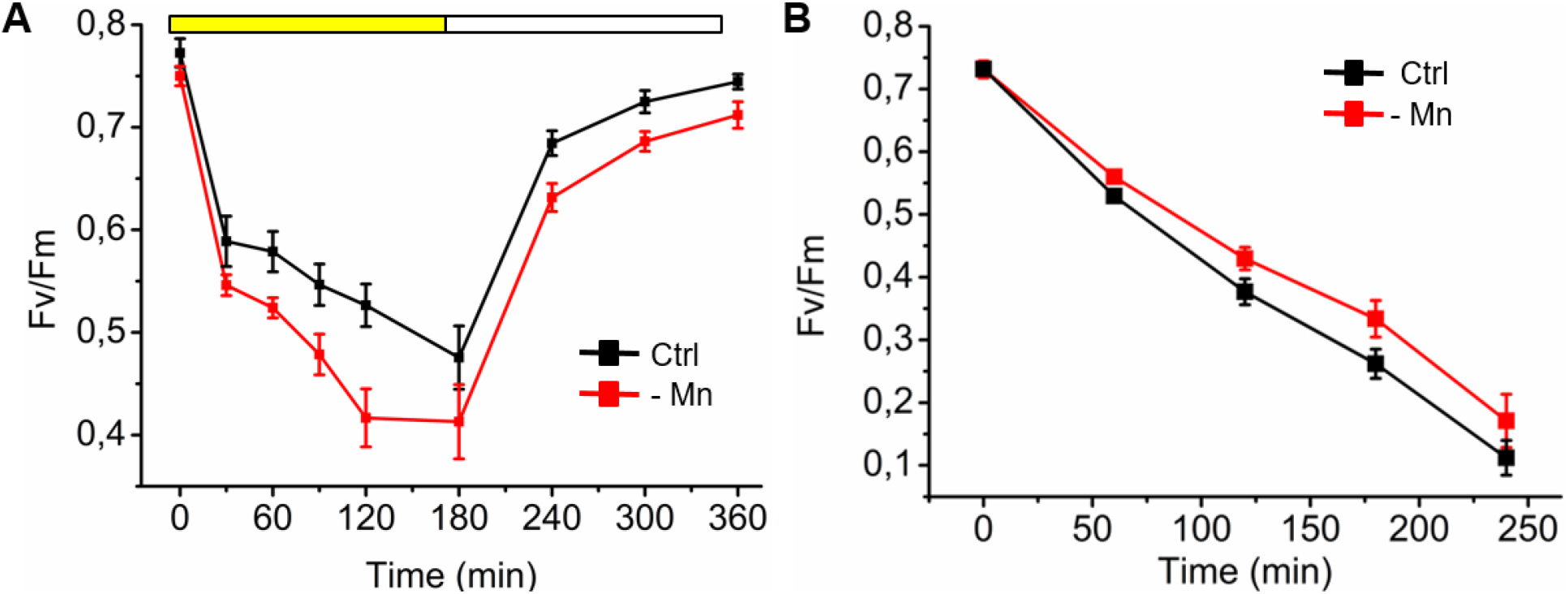
Photoinhibition in Mn deficiency in the absence and presence of lincomycin. **A:** Photoinhibition without lincomycin was measured on starch cultivated thalli with (black symbols) or without Mn (red symbols). Fv/Fm was followed on plants exposed to a strong light (800 μmol photons m^-2^s^-1^) for 3h (yellow bar) and during the recovery at room light (white bar). **B:** Fv/Fm was recorded during photoinhibition with lincomycin on starch cultivated plants with (black) or without Mn (red) for 4h (N=3, biological replicates).

The higher PSI fluorescence at 77K, the higher NPQ and the protection of PSII against photoinhibition in plants cultivated on Starch-Mn may indicate that cyclic electron transport is favoured under these conditions. Absorption changes of P700 show that P700 is more slowly oxidized, both in red actinic and in far-red light, in plants cultivated on Starch-Mn indicating that more electrons are available in the electron transport chain (Fig. 8A). Fig. 8B shows less donor-side limitation of PSI over a whole range of light intensities in plants cultivated on Starch-Mn compared to those cultivated on Starch Control. These data further indicate a higher activity of cyclic electron flow under Mn-deficiency. Under both conditions, acceptor side limitation was not observed because Marchantia possesses flavodiiron proteins (Shimakawa et al., 2021).

**Figure 8:**
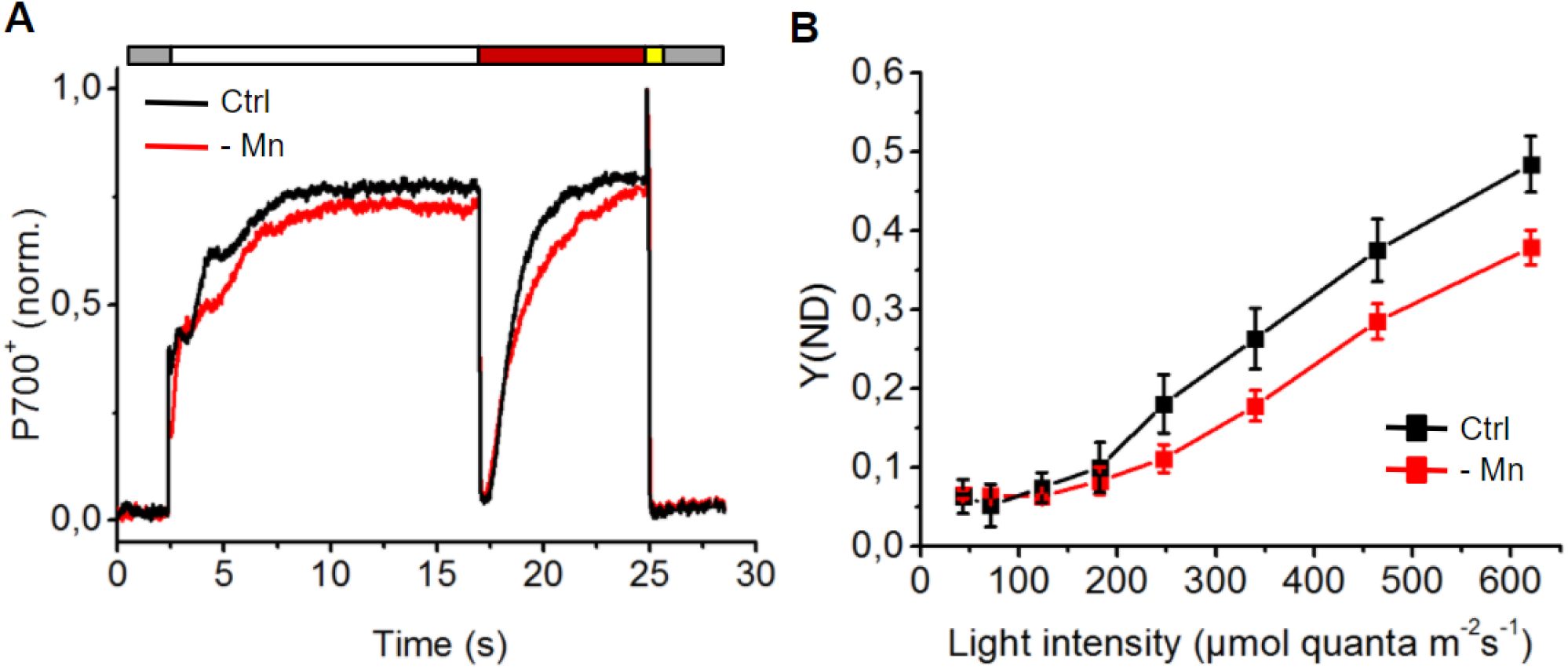
Activity of photosystem I in Mn deficiency. **A:** Normalized P700^+^ signal was obtained on thalli cultivated on Starch (+Mn, black; -Mn, red). The redox state of the PSI primary donor P700 was monitored through the changes in absorbance at 830 versus 875 nm. Leaves were kept in the dark for 5 min prior to the measurements. Thalli were exposed to actinic light for 15s (white bar) and to far-red light until reaching a plateau (red bar). Then, a saturating flash was applied (yellow bar) and the decay was observed in the dark (grey bar). **B:** PSI donor-side limitation Y(ND) based on saturating pulse analyses. Following the initial determination of maximal oxidation of P700, actinic light of the indicated intensities was given for 180 s.

## Discussion

In this study, we established growth conditions for manganese excess and deficiency in *Marchantia polymorpha*. Marchantia is able to absorb up to 25 mg Mn g DW^-1^ (Fig. 1b) which is higher than the hyperaccumulator *Polygonum lapathifolium* which contains approximately 18 mg Mn g DW^-1^ in its aerial parts (Liu et al., 2016). However, Marchantia suffers under the 200X conditions, and thalli only survive for a few weeks. Therefore, Marchantia does not qualify as hyperaccumulator despite its high Mn uptake capacity. Mn excess conditions induced an increase in metabolites involved in heavy metal detoxification and protection against oxidative stress as well as an increase in the SOD and PRX activity while catalase activity decreased (Table 1). A higher SOD activity and a large increase in N-methylalanine was observed also already in the 30X condition, showing that this Mn concentration is already stressful for the plant. Increase in SOD activity could reflect a higher amount of MnSOD present under Mn excess conditions. In angiosperms, Mn excess is known to increase the content of amino acids, organic acids or phenolic compounds, which are mainly involved in heavy metals stress tolerance (Alejandro et al., 2020; Singh et al., 2021), but this was not the case in Marchantia (Fig. 2). However, an accumulation of tryptophan was observed, which could be involved in the synthesis of tannins as has been reported under Mn excess (Alejandro et al., 2020). On the contrary, N-methylalanine seems to play a central role in the response to Mn excess in Marchantia. N-methylalanine is close to beta-alanine known to protect angiosperms against various stresses including heavy metal stress (Klapheck et al., 1988). N-methylalanine is most likely generated by oxidation of spermidine. It seems possible that the polyamine synthesis pathways in Marchantia are slightly different from those in Angiosperms. Further experiments should be conducted to determine whether N-methylalanine arises indeed from spermidine or putrescine oxidation (also increased under our excess conditions) and whether it is able to directly chelate Mn as it has been shown *in vitro* for Co(III) (Masahiko and Sado, 1968). Inside the chloroplast, most common molecules such as tryptophan, threonine and citrulline, could play the role of protection against heavy metal directly by chelating Mn or interacting with metal transporters, or through mitigating oxidative stresses, allowing Marchantia to maintain a certain level of photosynthetic activity and to survive in these drastic conditions. Mn excess affected the ratio between Mn and other metals such as Fe and Mg inside the thalli and the chloroplasts. It is known that divalent cation transporters are not completely selective and are capable of transporting several metals depending on their abundance in the medium. In fact, transporters such as IRT1 (Iron Regulated Transporter 1), known as an iron transporter have been shown to be able to transport also zinc, manganese, cobalt and cadmium (Barberon et al., 2011; Vert et al., 2002). It has also been shown that an excess of manganese in the soil prevents the absorption and translocation of other elements including iron, magnesium and phosphorus (Millaleo et al., 2010; Ducic et al., 2005; Lei et al., 2007) and can thereby lead to Fe deficiency. However, Marchantia grown at 200X does not show alterations in the Fe and Mg content inside the thalli (Figs 1B, 2A). Nevertheless, an alteration in the Mn/Fe ratio may induce Fe deficiency through competitive binding. Fe deficiency is known to affect primarily PSI that is rich in 4Fe4S clusters. As shown in Fig. 5B, PSI content is lowered in comparison to PSII in 200X. A similar reduction of PSI upon exposure to Mn excess has been reported for Arabidopsis (Millaleo et al., 2013). Aside PSI, the water-splitting activity of PSII seems also to be affected in the 200X condition as seen by the appearance of the K phase in fluorescence induction curves (Fig. 5C) and the decrease in PSII activity (Table 2). A toxic effect of Mn excess on PSII activity is in accordance with data on higher plants (Liang et al., 2019). 200X Mn negatively affected not only the photosynthetic apparatus but also led to a reduction in chloroplast size and an alteration of the well-separated grana stack and stroma lamellae distribution (Figs 3,4).

Transfer of thalli from agar to starch plates allowed to establish Mn deficiency conditions. Growth on starch affects the physiology of *M. polymorpha*. Comparison between the metabolite data of the two starch conditions with the Agar Control shows a severe reduction of the content of nine amino acids that are essential for plant growth (SI Table 2). Growth on starch allowed lowering drastically the Mn content in the thalli to 21 μg Mn g DW^-1^. According to Mengel and Kirkby (1987), the minimum Mn content required for growth of higher plants is about 20 to 40 mg Mn kg DW^-1^, and most higher plants usually contain 30 to 500 mg Mn kg DW^-1^. Mn deficiency led to a decrease in Fv/Fm, a change in the ratio of the activity PSI/PSII (Table 2) and less PSII content relative to PSI according to 77K fluorescence (Fig. 5B). In higher plants, Mn deficiency lowers PSII activity while PSI activity remains unaffected (Homann, 1967). Furthermore, the chloroplast size was decreased (Fig. 3) and the structural organization of the thylakoids was affected (Fig. 4). It is known that Mn deficiency leads to disorganization of the thylakoid membrane in higher plants (Mercer et al., 1962; Homann, 1967). In *M. polymorpha*, the smaller compartmentation of the grana stacks seems to be partly responsible for the changes in electron transport. In addition to a change in the stoichiometry between active PSI and active PSII, a different distribution of photosynthetic complexes may favour cyclic electron flow around PSI by allowing the formation of supercomplexes required for cyclic electron flow (Iwai et al., 2010). As shown in Fig. 8, thalli grown on Starch-Mn show less limitation of electron donation to P700^+^ than those on Starch Control indicating a stimulation of cyclic electron flow under Mn deprivation. Cyclic electron flow is known to be induced under specific physiological conditions like anaerobiosis in *Chlamydomonas reinhardtii*. Changes in the organization of PSI complexes was also observed under Mn limitation in cyanobacteria (Salomon and Keren, 2011). Increased cyclic electron flow generates a higher proton gradient across the thylakoid membrane and thereby an increase in NPQ (Fig. 5c). A higher NPQ protects photosystem II against photoinhibition as seen when thalli grown on Starch-Mn were exposed to high light in the presence of lincomycin (Fig. 7b) while they were more susceptible to photoinhibition in the absence of lincomycin (Fig. 7a) due to a slowdown of PSII repair. A similar slowdown of PSII repair under Mn deficiency has been observed previously in the Mn transporter mutant *nramp3nramp4* in Arabidopsis (Lanquar et al., 2010). The repair of PSII may be slowed down because of the higher ROS level in Starch-Mn (Table 2) that affects the synthesis of the D1 protein (Nishiyama *et al*., 2011) or by the lower availability of Mn for photoactivation, i.e., the light-dependent assembly of the Mn_4_CaO_5_ cluster. Future work is needed to explore the link between changes in the organization of the thylakoid membrane and stimulation of cyclic electron flow under Mn deficiency.

## Material and Methods

### Plant growth conditions

Gemmae from *Marchantia polymorpha*, Takaragaike (Tak-1) accession (male) were asexually cultured on ½ Gamborg’s GB5 1% Agar medium (Gamborg et al.,1968) for two weeks under a light-dark cycle of 16 h-light, 22°C, 120 μmol quanta m^-2^s^-1^, white fluorescent lamp / 8 h-dark, 20°C. For Mn excess condition, young thalli were transferred on ½ Gamborg’s GB5 1% Agar medium containing 33 μM (control), 0.5 mM (15X), 1 mM (30X), 2 mM (60X), 5 mM (150X) or 6.5 mM (200X) MnCl_2_ for one week before measurements. For Mn deficiency, young thalli were transferred on 1/2 Gamborg’s GB5 6% starch medium with or without addition of 33 μM MnCl_2_ for one week before performing measurements.

### Chloroplast and thylakoid isolation

10 g thalli were collected and ground in 100 ml of GR buffer (50 mM Hepes-KOH pH 7.5, 0.33 M sorbitol, 1 mM MgCl_2_, 1 mM MnCl_2_, 2 mM EDTA, 5 mM Na-ascorbate) with a spatula of BSA. Slurry was filtered through two layers of Miracloth and centrifuged at 1200*g* (4°C) for 7 min. Pellets were resuspended in 2 mL of GR buffer. The resuspended pellet was transferred on a percoll gradient (15 ml of 30% percoll/GR buffer and 10 ml of 70% Percoll/GR buffer), centrifuged at 7000*g* (4°C) for 17 min (no brake). Intact chloroplasts were collected and washed in 25 ml of GR buffer; centrifugation 1500*g* (4°C) for 5 min. For thylakoids, the same procedure was used without the percoll gradient.

### Metal quantification

Thalli were collected and dried at 70°C for 2 days before being weighted. 1 mL of nitric acid was added to 1 mg plant material and after complete dissolution samples. For intact chloroplasts, the chlorophyll concentration was determined before 0.5 mL nitric acid was added (250 μL sample with 100 μg chl mL^-1^). After 10-fold dilution in trace-metal-free water, the metal content of the samples was determined by atomic emission spectroscopy using a MP AES 1200 spectrometer (Agilent, USA).

### Super-resolution microscopy

For each sample, a small quantity of thallus was placed in a sandwich of two glass coverslips (22 mm diameter, Paul Marienfeld GmbH & Co. KG), sealed and placed inside an in-house built sample holder. The sample holder was fixed to a nano-positioning system (P-733.2CD, P-725.4CD, E-725.3CD, Physikinstrumente) coupled to an inverted microscope (Ti-U, Nikon) equipped with a CFI Apochromat Lambda 100x oil immersion objective. Room-temperature fluorescence emission was recorded by an iXon ULTRA 897 camera (Andor Technology Ltd., Belfast) coupled to the microscope with a custom-made Optomask (Cairn research Ltd, Faversham). Total chlorophyll fluorescence emission was isolated using a dichroic mirror/emission filter doublet (59022BS/ET655LP, Chroma Technology Corporation). Epifluorescence images were recorded using an excitation provided by a plasma light source (HPLS245 Thorlabs, Inc.) and an excitation filter (MF469-35 Thorlabs, Inc.). The excitation source for the laser-scanning measurements was a 445 nm emitting laser (OBIS LX, Coherent Inc.). Laser-scanning and final image reconstruction (resolution (*d*) *d*_xy_=126 nm, *d*_z_=320 nm), was as previously described (Streckaite, 2021) with a dwell time of 10-30 ms, 60 nm X/Y-scan steps and 300 nm Z-scan steps.

### Pigment analysis

Thalli were weighed and incubated in 100% acetone for 16 h in the dark. Pigment extract was diluted to 80% acetone before measurement. Chlorophyll *a* and chlorophyll *b* contents were calculated according to Arnon, 1949.

### 77K Chlorophyll fluorescence measurements

Fluorescence spectra of intact chloroplasts diluted in GR buffer were measured with a Carry Eclipse fluorimeter; excitation wavelength: 430 nm. The intensity was normalized to the intensity of the PSI emission.

### Chlorophyll fluorescence analysis at room temperature

Chlorophyll fluorescence analysis was performed on 5 min dark-adapted thalli using a Dual-PAM-100 fluorimeter (Walz, Effeltrich, Germany). Fm and Fm’ were determined using saturating flashes (10,000 μmol quanta m^-2^s^-1^, duration 300 ms). Fo, minimum and Fm, maximum fluorescence in a dark-adapted sample; Fm’, maximum fluorescence and F’, fluorescence emission from a light-adapted sample. Induction and recovery curves were measured using actinic light of 50 μmol quanta m^-2^s^-1^. F_v_/F_m_ = (F_m_-F_0_)/F_m_, qN = (F_m_-F_m_’)/(F_m_-F_0_’) and qP = (F_m_’-F)/(F_m_’-F_0_’). For induction curves (OJIP) one saturating flash was given.

### Photoinhibition

Thalli were placed on wet filter paper and illuminated with white light (800 μmol quanta m^-2^s^-1^); LED panel SL3500 (Photon Systems Instrument, Drasov, Czech Republic). For recovery, samples were placed in low light (10 μmol quanta m^-2^s^-1^). When indicated, thalli were incubated in lincomycin (1 g L^-1^) for 4 h prior to the photoinhibition treatment. Fv/Fm was measured using an Imaging-PAM (Walz, Effeltrich, Germany).

### P700 measurements

P700 absorbance was measured using a Dual-PAM-100 fluorometer. Near-infrared measuring lights (830 and 870 nm) were applied to measure the transmittance of oxidized P700. Prior to the measurements, the plants were kept in the light in the growth chamber so that the Calvin-Benson cycle enzymes were active. Five minutes dark-adapted thalli were exposed to actinic light followed by far-red light. Then, a saturating flash was given at the end of the far-red light period. To determine quantum yields of PSI donor (Y(ND)) and acceptor side-limitations (Y(NA)) saturating pulse analysis was used (Klughammer and Schreiber, 1994). Each actinic light intensity was applied for 180 s before determining Y(ND) and Y(NA).

### Antioxidant enzymes activity

Thalli were harvested and ground in 50 mM HEPES buffer pH 6.5. The slurry was filtered through one layer of miracloth, centrifuged for 5 min at 10.000*g* (4°C). Protein concentration in the crude extract was determined by amido black since the supernatant contained pigments. 10 μg mL^-1^ protein was used for the enzymatic tests. Guaiacol peroxidase activity was determined spectrophotometrically by measuring the oxidation of guaiacol to tetraguaiacol at 470 nm (ε: 26.6 mM^-1^cm^-1^). The reaction mixture contained 50 mM NaH_2_PO_4_/Na_2_HPO_4_ pH 7.5, 3 mM H_2_O_2_, 0.01% (v/v) guaiacol. Catalase activity was measured polarographically at 20 °C with a Clark-type electrode in 50 mM Tris buffer pH 8.5 and 1 mM H_2_O_2_ as substrate. Superoxide dismutase activity was measured using a solution of 500 μM Xanthine, 20 mM HEPES buffer pH 7; 0.2 U mL^-1^ xanthine oxidase and 100 μM XTT (Na,3’-(1-(phenylaminocarbonyl)-3,4-tetrazolium)-bis-(4-methoxy-6-nitro) benzene sulfonic acid hydrate) as substrate. The kinetics of superoxide production were measured as increase in absorbance at 470 nm, and the SOD activity was determined by following the inhibition of the superoxide production after addition of the crude extract. For ^•^OH detection, 15 mg thalli were incubated for 1 h in 3 ml 10 mM phosphate buffer, pH 6.0, 50 mM N-tert-butyl-α-(4-pyridyl)nitrone N’-oxide (4-POBN) and 4% ethanol (Heyno et al., 2008).

### Metabolite Analysis

GC-MS profiling of metabolites was performed on thalli cultured on Agar Control, 30X and 200X conditions and thalli grown on Starch Control and Starch-Mn. Dried and grounded samples (5 mg of dry weight) were incubated in 1 mL of H_2_O/ACN/isopropanol (2/3/3) with 4 mg l^-1^ of ribitol for 10 min at 4°C with shaking at 250g in an Eppendorf Thermomixer. Insoluble material was removed by centrifugation at 20 400g for 10 min. 700 μl of supernatant were recovered, 70 μl of a mix of H_2_O/MeOH/isopropanol (2/5/2) with 0.3 g L^-1^ of myristic acid d27 was added as an internal standard for retention time locking. Aliquots of each extract (100 μL) extracts were dried for 4 h at 30°C in a Speed-Vac and stored at −80°C. After thawing, samples were dried again in a Speed-Vac for 1 h at 30°C before adding 10 μL of 20 mg mL^-1^ methoxyamine in pyridine to the samples. The reaction was performed for 90 min at 30°C under continuous shaking in an Eppendorf thermomixer. A volume of 90 μL of N-methyl-N-trimethylsilyl-trifluoroacetamide was then added and the reaction continued for 30 min at 37°C. After cooling, 80 μL were transferred to an Agilent vial for injection. Four hour after derivatization, 1 μL sample was injected in splitless mode on an Agilent 7890B gas chromatograph coupled to an Agilent 5977A mass spectrometer (column: RESTEK RXI 5SIL MS 30MX0.25MMX0.25UM). An injection in split mode with a ratio of 1:30 was systematically performed for quantification of saturated compounds. Raw Agilent data files were analysed with AMDIS http://chemdata.nist.gov/mass-spc/amdis/. The Agilent Fiehn GC-MS Metabolomics RTL Library was employed for metabolite identifications. Peak areas were determined with the Masshunter Quantitative Analysis (Agilent) in splitless and split 30 modes. Peak areas were normalized to ribitol and dry weight. Metabolite contents are expressed in arbitrary units (semi-quantitative determination).

## Acknowledgements

This work was supported by the Labex Saclay Plant Sciences-SPS (ANR-17-EUR-0007), and the French Infrastructure for Integrated Structural Biology (FRISBI; grant number ANR-10-INSB-05). M.M. is supported by a CEA PhD fellowship.

## Author Contribution

M.M, and A.K-L. designed the project. M.M, T.H., F.G., B.G., A.G., S.T. and A.K-L. performed the experiments and analysed the data. M.M. and A.K-L. wrote the initial version of the manuscript that was read and revised by all authors.

## Data availability

Data will be made available on demand.

## Supporting Information

**Table S1** List of 96 metabolites identified by GC-MS in *Marchantia polymorpha*

**Table S2** Metabolites significantly decreased in Starch condition compared to Agar Control in *Marchantia polymorpha*

**Figure S1** Chlorophyll fluorescence measurements at room temperature for excess conditions

**Figure S2** Photoinhibition experiment with and without lincomycin for excess conditions

